# A *Drosophila* metainflammation-blood tumor model links aspirin-triggered eicosanoid-like mediators to immune signaling

**DOI:** 10.1101/624197

**Authors:** Silvio Panettieri, Indira Paddibhatla, Jennifer Chou, Roma Rajwani, Rebecca S. Moore, Tamara Goncharuk, George John, Shubha Govind

## Abstract

Accumulating data from epidemiologic studies are linking aspirin’s use to a decline in chronic and metabolic inflammation that underlies many human diseases including some cancers. Aspirin reduces cyclooxygenase-mediated pro-inflammatory prostaglandins and promotes the production of pro-resolution molecules. Aspirin also triggers the production of anti-inflammatory electrophilic mono-oxygenated lipid mediators implicated in human pathologies. With the goal of developing a model system for studying the mechanisms of aspirin in reducing inflammation, we investigated aspirin’s effects in fruit fly models of chronic inflammation. Ectopic Toll/NF-κB and JAK/STAT signaling in *D. melanogaster* results in an overproliferation of hematopoietic blood progenitors coupled with metabolic inflammation in adipocytes. We report that, like mammals, flies are sensitive to aspirin treatment and it modulates the Toll-NF-κB axis. Aspirin-treated mutants simultaneously experience reduction in metabolic inflammation, mitosis, ectopic immune signaling, and macrophage infiltration. Moreover, flies synthesize 13-HODE, and aspirin triggers 13-EFOX-L2 production in mutants. In such flies with ectopic immune signaling, providing 13-HODE’s precursor linoleic acid or performing targeted knockdown of transcription factor STAT in inflammatory blood cells boosts 13-EFOX-L2 levels while decreasing metabolic inflammation. Thus, hematopoietic cells regulate metabolic inflammation in flies, and their effects can be reversed by pharmaceutical or dietary intervention, suggesting deep phylogenetic conservation in animals’ ability to resolve systemic inflammation and repair tissue damage. This model system brings the power of *Drosophila* genetics to bear on immuno-metabolic mechanisms that boost systemic health and healing, with the potential to identify new targets for the treatment of chronic diseases in humans.

## Introduction

As part of an animal’s innate immune system, acute inflammation targets parasites and microbes and repairs injured tissues. Circulating macrophages and neutrophils migrate to sites of infection or injury where they initiate acute inflammation either to phagocytose and clear invading organisms, or to sequester parasite or cellular debris. Mammals and *Drosophila* rely on the archetypal pro-inflammatory NF-κB signaling pathway to activate the inflammation response (1). Canonical NF-κB signaling employs Toll/Toll-like receptors (TLRs), which, when bound by pathogen-derived ligand in mammals, mediate the release (from their I-κB inhibitors) and nuclear import of NF-κB. NF-κB activates pro-inflammatory pattern of gene expression (1,2). Pathogen receptors in flies also activate Toll/NF-κB signaling for host defense (3-5).

A key target of pro-inflammatory NF-κB activation in mammals that has garnered much attention is the prostaglandin-endoperoxide synthase 2/cyclooxygenase-2 (*PTGS2/COX-2*) promoter (6-8). COX-2 catalyzes the rate-limiting step in the conversion of the polyunsaturated C20:4 ω-6 fatty acid (FA), arachidonic acid (AA), to pro-/anti-inflammatory prostanoids (9). AA, an important component of the plasma membrane’s phospholipids, is derived from dietary omega-6 FA, linoleic acid (LA). Aspirin (acetylsalicylic acid, ASA) blocks the pro-inflammatory effects of prostanoids by acetylating and inhibiting COX-1 and -2 activities (10). NF-κB-mediated induction of *COX-2* integrates eicosanoid-mediated signaling with NF-κB signaling. Toll/NF-κB signaling also responds to free fatty acid levels in the blood. For example, saturated (but not unsaturated) fatty acids activate TLR4 and its downstream mediators, and NF-κB p65 activation triggers *COX-2* expression (11,12). Since Toll receptors sense both pathogens and nutrients in mammals, diet and energy balance are closely integrated with innate immunity and inflammation (2,13,14).

A second biochemical effect of aspirin was identified in the resolution of acute inflammation, which occurs due to its ability to acetylate COX-2. Many studies have identified a new class of specialized pro-resolving mediators (SPMs) formed in the presence of acetylated COX-2. Derived from C20 or C22 ω-3 and ω-6 FAs, SPMs include lipoxins, resolvins, protectins, and maresins. SPMs boost efferocytosis, promote phagocytosis of microbes, and limit neutrophil-mediated tissue damage (15). In human cell cultures, acetylated COX-2 also triggers the production of a family of electrophilic mono-oxygenated (EFOX) lipid mediators, including 17- and 13-EFOX, derived from ω-3 C22:6 docasahexanoic acid (DHA) (16). While the existence and anti-inflammatory effects of EFOX in intact organisms have not been demonstrated yet, results of these cell culture studies point to the dual anti-inflammatory and pro-resolution roles of aspirin and suggest an explanation for how dietary supplementation of ω-3 FAs or aspirin stalls inflammation and supports well-being in humans (9,16). They also expose the diversity of eicosanoid and eicosanoid-like lipid mediators and the complexity of their cell-specific effects in maintaining normal organismic physiology. How are the pro- and anti-inflammatory effects of eicosanoid family members precisely integrated with metabolic and immune pathways *in vivo*? How are their effects unified from cellular to tissue and organ levels (e.g., from inflammatory blood cells to adipose tissue)? Because of their inherent diversity and cellular complexity, vertebrate models do not lend themselves to straightforward resolution of such questions.

To develop a simpler model for systematically addressing these issues, we used *D. melanogaster* models with proven hallmarks of chronic inflammation (17-20). As in mammals, innate immunity in *Drosophila* also relies on NF-κB signaling (3-5). Furthermore, systemic acute inflammation is induced by parasitic wasp infection. Injection of an egg into a fly larva activates inflammatory phagocytic macrophages (plasmatocytes) and their non-phagocytic derivatives (lamellocytes), which migrate and adhere to the wasp egg, blocking its development. This process relies on: (a) Spätzle (Spz), the pro-mitotic and pro-inflammatory cytokine and Toll ligand; (b) Spätzle Processing Enzyme (SPE); and (c) NF-κB transcription factors, Dorsal and Dif (18). The canonical Toll/NF-κB target gene Drosomycin (*Drs*) is activated after wasp infection, as is *cactus*, the gene coding for the I-κB inhibitor of Dorsal/Dif (18). In another parallel with mammalian immunity, the negative regulator Ubc9 (the E2 SUMO-conjugating enzyme) maintains immune homeostasis in *Drosophil*a. As a consequence of persistent NF-κB signaling in *cactus* and *Ubc9* mutants, small inflammatory blood tumors, made of macrophages and lamellocytes, appear in the larval body cavity. Overexpression of SPE or Spz in blood cells of wild type animals induces similar tumors while reduction of *spz* lowers inflammation in *Ubc9* mutants with concomitant reduction of tumors (18).

*Ubc9* mutants also exhibit significant loss of integrity in the basement membrane surrounding the liver-like fat body; collagen distribution is discontinuous and many macrophages infiltrate (i.e., adhere to) the diseased tissue. Wild type fat body is surrounded by a uniform and continuous deposit of collagen and has few, if any, macrophages adhering to it (18,21). Thus, akin to mammals, tissue damage in flies accompanies systemic inflammation (1,2,18,22).

Flies not only possess these conserved pro-inflammatory immune pathways for the activation of inflammation, but also react to metabolic stress in ways similar to humans. In response to a high-sugar diet (but not to excess dietary fat or protein), *Drosophila* develop signs of metabolic inflammation (or metainflammation) where cellular and molecular phenotypes resemble the pathophysiology of type-2 diabetes mellitus (2,23). High-sugar-fed larvae exhibit an altered metabolism and the cells of their fat body organ display larger-sized lipid droplets (LDs, which store and mobilize fat in animal cells (23)). Therefore, mechanistically, sugar-induced metabolic dysregulation in flies resembles insulin resistance in humans (13,21,23).

Given that flies exhibit hallmarks of parasite-induced inflammatory and diet-induced metainflammatory responses, we asked if these phenotypes are sensitive to aspirin, and if so, how aspirin might activate and coordinate animals’ intrinsic healing mechanisms. While flies lack FAs longer than C18 (AA, DHA, etc.) (24) and eicosanoids have not been reported from *Drosophila*, prostaglandin administration rescues the loss-of-function of a human *COX-1*-like fly gene (25), suggesting that flies possess primordial eicosanoid-based mechanisms through which aspirin might act. In addition, C20 polyunsaturated fatty acid (PUFA)-fed flies generate hydroxyeicosatetraenoic acid (13-HETE) (26), an AA metabolite, suggesting that flies express enzymes for eicosanoid synthesis. Using computational methods, we recently showed that functional enzymes for eicosanoid biosynthesis are encoded in the fly genome. This analysis revealed the existence of two other COX-like enzymes in addition to Pxt (27). We therefore hypothesized that inflammation readouts in flies may be sensitive to aspirin and flies might use simpler FA precursors to produce eicosanoid-like molecules.

Here we show aspirin’s dose-dependent and molecular-, cell-, and organ-specific effects with striking parallels to its effects in mammals. We also show that flies produce 13-HODE, and aspirin increases its levels, while also triggering the production of its oxidized analog 13-oxo-ODE (called 13-EFOX-L2 here). These results demonstrate that members of the EFOX family of oxylipids are produced in intact organisms and their high levels correlate with a reduction in inflammation. Aspirin’s efficacy in resolving inflammation and repairing tissue damage in fruit flies suggests genetic and biochemical conservation underlying these processes across the animal kingdom. Unbiased genetic screens in *Drosophila* that report the links between inflammation, diet, and drug action will have translational significance in identifying new therapies for inflammation-based human diseases.

## Results

Wild type *Canton S* flies tolerate a wide range of orally administered aspirin. At 5 mM aspirin, 65% of the animals eclosed to adulthood while 42% of pupae became adults at 55 mM. In experiments described below, 0.5 nM to 1 mM aspirin was added to fly food (see SI for additional details). Aspirin uptake was confirmed by covalently conjugating it to Rhodamine B. Dissected hematopoietic and fat body cells did not retain the unconjugated dye *ex vivo*, but Rhodamine B-aspirin (Rh-ASA) localized to the perinuclear regions in both cell types (SI Fig. S1).

### Aspirin inhibits hematopoietic overgrowth in *Ubc9* mutants

We introduced 0.5 nM or 5.5 μM aspirin to the diet of developmentally synchronized cultures of heterozygote and inflammatory tumor-producing homozygous *Ubc9* mutant larvae, and examined its effects on tumor abundance (penetrance) and sizes (tumor projection area or expressivity), gene expression, and LD sizes in the fat body. Aspirin (0.5 nM and 5.5 μM) treatment resulted in a remarkable increase in tumor-free animals (0-2 tumors/animal; Fig. 1A, B, D). Treatment at both concentrations reduced tumors in animals with three or more tumors somewhat differently, although at both concentrations, tumor projection area shrank by more than 60% (Fig. 1C, E).

**Fig. 1.**
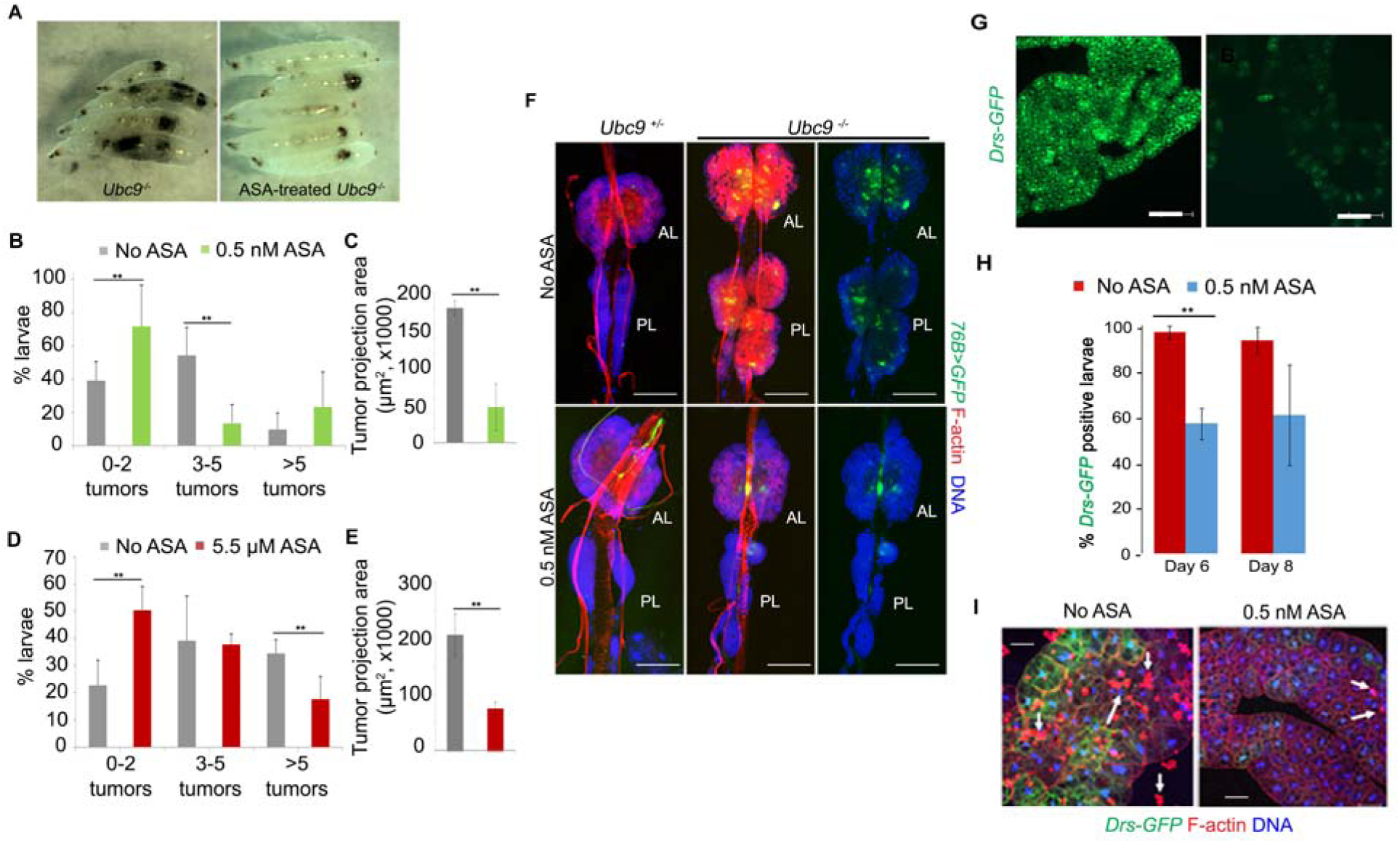
Anti-inflammatory effects of aspirin in the *Ubc9* model of chronic inflammation. **(A)** Untreated (left) and 0.5 nM aspirin-treated *Ubc9* 8-day-old larvae showing clearance of some but not all tumors. **(B-E)** Tumor penetrance (B, D) and tumor size (average of measured projection areas of all tumors (C, E) in 8-day old *Ubc9* larvae. For tumor penetrance (B and D), the proportion of mutants with 0-2 tumors to 5 or more tumors is shown. Data are from three biological replicates. The number of animals for each replicate and treatment ranged from 5-20. Tumor penetrance and tumor size data were obtained from the same animals. Bars represent standard deviation and ** represent *p* value < 0.05. **(F)** Aspirin shrinks the expanded hematopoietic population in *Ubc9* mutants. Anterior and posterior lymph gland lobes from untreated or 0.5 nM aspirin-treated control or mutant animals. The progenitor population is marked with *76>GFP*. Cells are stained with Phalloidin to visualize actin-rich lamellocytes and dorsal vessel; Hoechst identified nuclei. AL, anterior lobes, PL, posterior lobes. Right panels do not show the red signal in mutant glands (Scale bar: 100 μm). **(G, H)** Aspirin administration attenuates *Drs-GFP* reporter expression in the *Ubc9* fat body. **(G)** Fat body tissue from a 6-day-old *Ubc9* larva after 0.5 nM aspirin administration shows significant reduction in the *Drs-GFP* expression (right) compared to fat body tissue from an untreated (left) *Ubc9* larva. Scale bar = 100 μm. **(H)** Percentage of larvae with *Drs-GFP* expression. Entire fat body was dissected to study GFP expression. A fat body was considered GFP-positive if at least 10 cells expressed GFP in the entire organ. The bars represent mean ± standard error from three biological replicates. At day 6, 96.7 ± 3% (untreated) and 56.9 ± 16% (0.5 nM aspirin-treated) larvae were GFP-positive. At the 8-day time point, 93.3 ± 7% (untreated) and 60.7 ± 23% (0.5 nM aspirin-treated) larvae were GFP-positive. The number of 6-day old larvae per replicate ranged from 8 to 19; the number of 8-day old larvae used per replicate ranged from 5 to 18 (few 8-day old animals survive). *p* = 0.03 for 6-day old larvae; *p* = 0.12 in 8-day old larvae; student t-test was applied. **(I)** *Drs-GFP* fat bodies from untreated (left) and aspirin-treated (right) 8-day-old *Ubc9* larvae show simultaneous reduction in blood cell infiltration (Rhodamine-Phalloidin; arrows) and *GFP* expression. (Scale bar: 50 μm).

We have previously shown that *Ubc9* tumors are derived from overproliferating lymph gland lobes (25). Reduction in tumor penetrance by aspirin treatment (Fig. 1A-E) was also reflected in the shrinkage of lymph gland lobes and in the restoration of lymph gland morphology (Fig. 1F). Relative to heterozygous lymph glands, *76B>GFP-*marked mutant *Ubc9* lymph glands have an expanded GFP-positive progenitor population with many inflammatory lamellocytes (Fig. 1F) (28). Treatment with 0.5 nM aspirin reduced the progenitor and mature blood cell populations (Fig. 1F). The size of the lymph gland lobes was reduced: while the projection areas of anterior lobes did not differ significantly (*p* = 0.34), the first set of posterior lobes shrank in size (*p* = 0.005; 20,907.038 ± 1,838.236 µm^2^ in untreated versus 9,221.446 ± 3,238.399 µm^2^ in 0.5 nM aspirin treated animals (mean ± standard deviation); 3 biological replicates; 4-6 lobes per replicate). Thus, administration of aspirin restricted lymph gland overgrowth and tumor development.

### Aspirin restores regulation of Toll-NF-κB signaling in *Ubc9* fat body

To examine if constitutive signaling in *Ubc9* fat bodies (20) is rescued by aspirin, we studied *Drs-GFP* expression and noticed a significant decrease. At day 6, there was a ∼40% decrease in the percentage of *Drs-GFP* positive larvae (0.5 nM ASA; Fig. 1G, H). Thus, it appears that as in mammalian cells (29), aspirin inhibits chronic inflammation in flies by modulating the Toll-NF-κB axis. This decrease correlated with reduced infiltration of blood cells into the fat body adipocytes (Fig. 1I).

Immune responses in mammals can modulate insulin signaling and nutrient storage (30), and conversely, immune pathways are activated in response to nutrient stress (31). Given that the Toll/Dorsal pathway is constitutively active in *Ubc9* fat bodies (Fig. 1G-I), we wondered if LD sizes are affected in mutants and if aspirin might reverse these effects. In flies, the diabetes-like metabolic state is reflected in larger LDs in larval fat bodies of animals consuming high dietary glucose (23). *Ubc9* fat bodies produce larger-than-normal LDs without additional glucose, and a slight increase is evident even in heterozygous cells compared to *y w* control (Fig. S2A-C). Aspirin treatment reverses this trend (Fig. S2D) and the damaged mutant fat body morphologically appears intact when stained with Rh-Phalloidin (Fig. 1I; (18)). These results suggest that as in vertebrate models and in human trials where anti-inflammatory therapies restore metabolic function (31,32), aspirin treatment in flies also reduces metabolic stress.

### Toll-NF-κB signaling is hyperactive in *hopscotch*^*Tum-l*^ blood cells; aspirin is anti-mitotic, and improves organismal viability

To corroborate the effects of aspirin in another genetic background and understand how it promotes healing and tissue repair, we turned to the genetically uniform germline *hopscotch*^*Tum-l*^ (*hop*^*Tum-l*^ or simply *Tum-l*) mutant, where JAK-STAT signaling is hyperactive due to a dominant gain-of-function point mutation (19). In these mutants, hyperactive STAT activity expands the hematopoietic progenitor population (17,19,33) and lymph gland dysplasia leads to tumor development and partial lethality (19,34) (Fig. 2B-C). Microarray analysis revealed high expression of key Toll pathway genes (*SPE, spz, Toll*, and *Dif*) in *hop*^*Tum-l*^ mutants (Fig. 2D) (34).

**Fig. 2.**
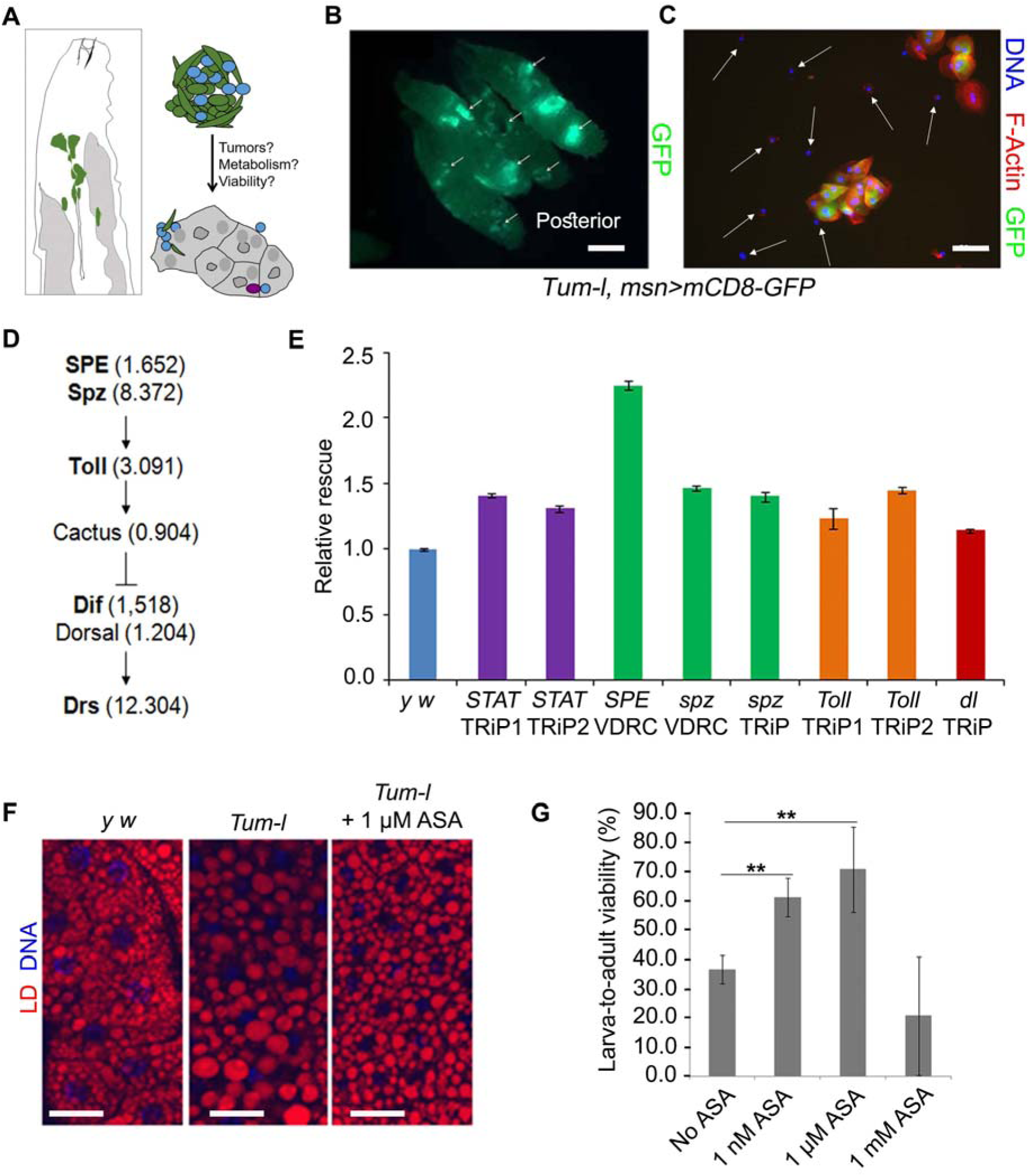
Aspirin is anti-inflammatory in *hop*^*Tum-l*^ larvae and improves viability. **(A)** Schematic showing *hop*^*Tum-l*^ *msn>mCD8-GFP* larva’s immune organs. An *msn-GAL4* transgene allows targeted manipulation of gene expression in progenitors of inflammatory lamellocytes. The effects of such manipulation on tumorogenesis, viability and metainflammation can thus be examined. **(B)** GFP-positive hematopoietic tumors (arrows) in 6-day old *hop*^*Tum-l*^ *msn>mCD8-GFP* whole larvae. (Scale bar: 1 mm). **(C)** An aggregate of mutant cells from 6-day old third instar larvae showing GFP-positive lamellocytes and GFP-negative macrophages (white arrows). (Scale bar: 50 μm). **(D)** Microarray data summarized from reference (34) performed on *hop*^*Tum-l*^ reveal that Toll pathway components are transcriptionally active. Significant increase (> 1.5-fold) is observed for *SPE, spz, Toll*, and *Dif* genes. **(E)** Relative viability of adult *hop*^*Tum-l*^ *msn>mCD8-GFP* males at 27°C with RNAi relative to viability of control *hop*^*Tum-l*^ males from an outcross of *hop*^*Tum-l*^*/FM7* females and *y w* males without RNAi. Bars represent mean ± standard error; *p* < 0.05 using student *t*-test for all pairwise comparisons across three biological replicates. The number of males scored for a given replicate ranged from 76 to 279. **(F)** Representative images of Nile red-stained larval fat body adipocytes from untreated *y w*, and untreated and 1 μM aspirin-treated *hop*^*Tum-l*^ *msn>mCD8-GFP* animals (scale bar, 30 μm). LD sizes are quantified in SI Fig. S4B. **(G)** Larval-to-adult viability of *hop*^*Tum-l*^ *msn>mCD8-GFP* animals upon systemic aspirin treatment (mean ± standard deviation is shown across four biological replicates; *n* > 38 larvae per replicate; ** *p* < 0.05; student *t*-test).

To determine if the expanded hematopoietic population is fundamental to chronic inflammation in *hop*^*Tum-l*^ mutants and if systemic effects compromise viability, we targeted the knockdown of *STAT* and *Toll* pathway genes (*SPE, spz, Toll*, and *dl*) in the expanding hematopoietic population (*hop*^*Tum-l*^ *msn>mCD8-GFP, RNAi)*. We compared the viability of “knockdown” animals with outcrossed *hop*^*Tum-l*^ mutants without knockdown. Viability of progeny from the *y w* outcross was no different if *white, ebony* or *GFP* RNAi was expressed (SI Fig. S3A). In contrast, we found that compared to the control *y w* outcross, *hop*^*Tum-l*^ mutants were more viable when pro-inflammatory STAT or NF-κB signaling was knocked down. The most significant gains were seen with *STAT, SPE*, and *spz* knockdown in blood cells (Fig. 2E). These results suggest a potential downstream (or cooperative) role of the Spz-Toll-Dorsal/Dif axis with STAT in *hop*^*Tum-l*^ blood cells. Aspirin treatment (1 μM) of *hop*^*Tum-l*^ animals reduced the mitotic index in circulating blood cells (SI Fig. S4A). These results confirm that, as such, aspirin’s effects are not limited to a specific genetic background, but instead its biological effects must be meditated by molecular pathways common to both mutants.

Like *Ubc9* mutants, adipocytes of the *hop*^*Tum-l*^ fat body form many large and distorted LDs without added dietary glucose (Fig. 2F; SI Fig. S4B). The mutant fat body loses integrity and intercellular adhesion; several portions of the organ detach from the main organ (SI Fig. S4C). However, administration of 1 µM aspirin restored a significant portion of the small LD population and there were fewer large LDs (Fig. 2F; SI Fig. S4B); aspirin also restored the overall morphology of the organ (SI Fig. S4D).

Encouraged by aspirin’s strong and consistent anti-tumor and anti-inflammatory effects at the molecular, cellular, and organ levels in two inflammation models, we asked if these effects might translate to benefits at the organismic level and if aspirin improves viability of *hop*^*Tum-l*^ animals. 1 nM and 1 μM aspirin improved the viability of *hop*^*Tum-l*^ animals to adulthood (Fig. 2G) and these gains paralleled viability increments in the targeted knockdown experiments (Fig. 2E). This correlation suggested that aspirin’s salutary effects are, in part, realized by blocking the systemic pro-inflammatory effects of blood cells. We hypothesized that, in addition to reducing mitosis and shrinking the pro-inflammatory blood cell population, aspirin might trigger oxylipid production whose anti-inflammatory effects may be realized in other parts of the body and possibly the entire organism. We tested this idea in biochemical experiments.

### Discovery of 13-HODE and 13-EFOX-L_2_ in fly larvae

Because *D. melanogaster* lack FAs longer than C18 (24), we searched for non-classical anti-inflammatory lipid mediators derived from unsaturated C18 precursors linoleic acid (LA) and alpha linolenic acid (ALA), present in the normal fly diet (24). Oxidized derivatives of LA and ALA (13-hydroxyoctadecadienoic acid (13-HODE) and 13-hydroxyoctadecatrienoic acid (13-HOTrE), respectively; structures in Fig. 3A and SI Fig. S5A) are bioactive and exert potent anti-inflammatory effects in mammals (9,15,16,32,35). In LC-MS/MS-based targeted lipidomics of larval extracts, we did not detect 13-HOTrE in wild type or mutant larvae, but discovered that untreated third instar wild type and *hop*^*Tum-l*^ larvae produce 13-HODE at levels comparable to mammalian cells (∼40 nM (Fig. 3B; Fig. S5B, D) (36)). Application of an enantiomer-specific antibody in ELISA experiments (Enzo Life Sciences) confirmed the presence of 13(S)-HODE in both isolated larval fat bodies and remaining organs of untreated larvae suggesting its wide occurence in the body. Chiral LC-MS/MS analysis resolved the R and S enantiomers of 13-HODE and showed that 13(S)-HODE was twice as abundant as 13(R)-HODE regardless of genetic background or aspirin treatment (SI Fig. S5D). This enantiomeric imbalance indicates that at least some of the oxidation product may be generated enzymatically. In mammalian models, the R and S enantiomers demonstrate pro-or anti-inflammatory activities.

**Fig. 3.**
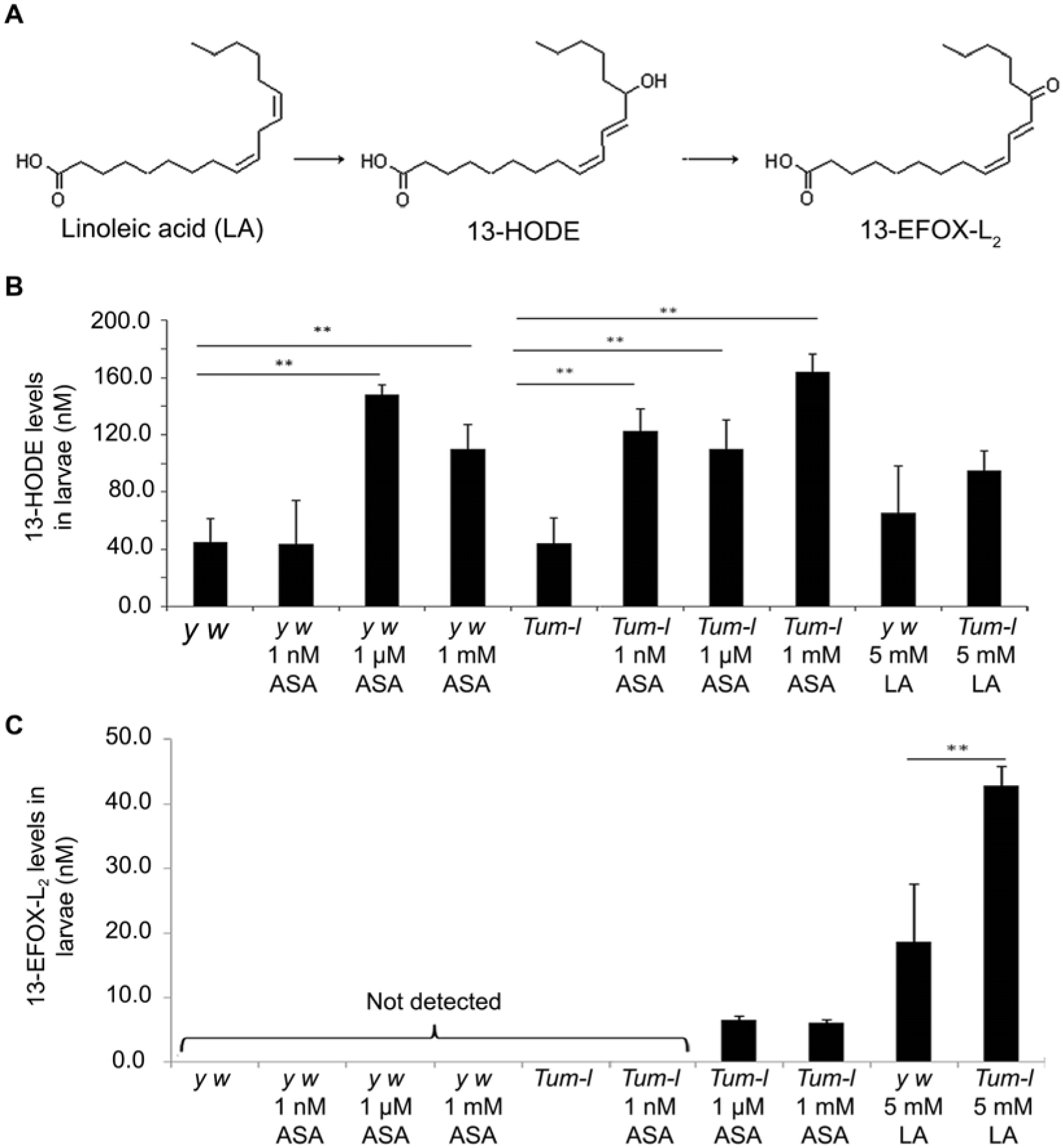
Aspirin (ASA)-triggered production of 13-HODE and 13-EFOX-L_2_. **(A)** LA is first converted into 13-HODE by COX-2 action, and then further oxidized to 13-13-oxo-ODE or EFOX-L_2_. This proposed scheme is based on ref. 7. **(B, C)** Quantification of 13-HODE (B) and 13-EFOX-L_2_ (C) levels in *y w* and *hop*^*Tum-l*^ animals without or with 1 nM, 1 μM, or 1 mM aspirin treatment (*n* > 200 larvae per measurement per compound, mean ± standard deviation across 3-5 biological replicates, \*\**p* < 0.05; one-way ANOVA was applied).

Quantitative LC-MS/MS further revealed that 13-HODE levels increase 3-fold in *y w* and *hop*^*Tum-l*^ larvae upon treatment with 1 μM and 1 mM aspirin (Fig. 3B). At 1 nM aspirin, however, a comparable 3-fold increase is observed only in the *hop*^*Tum-l*^ mutants but not in wild type animals (Fig. 3B), suggesting different response thresholds in these genetic backgrounds.

### 13-EFOX-L_2_ levels rise in response to aspirin and linoleic acid supplement

The hydroxyl group in 13-HODE, responsible for its chiral nature, can be further oxidized to generate the corresponding electrophilic derivative 13-EFOX-L_2_ according to the mechanism proposed by Groeger et al. for other PUFAs (9) (Fig. 3A). We detected 13-EFOX-L_2_ in only *hop*^*Tum-l*^ mutants at high aspirin concentrations (1 μM and 1 mM) but not in control *y w* animals (Fig. 3C; SI Fig. S5C). Appearance of 13-EFOX-L_2_ correlate with the rise in 13-HODE in response to aspirin treatment of mutants (Fig. 3B), supporting the idea that 13-HODE is a substrate for 13-EFOX-L_2_ synthesis *in vivo* (Fig. 3A).

The discovery of 13-HODE and 13-EFOX-L_2_, two LA-derived oxylipids in larval extracts prompted us to examine whether *hop*^*Tum-l*^ mutants would benefit from dietary addition of LA (13-HODE and 13-EFOX-L_2_ are chemically unstable for such experiments). 5 mM LA treatment increased *hop*^*Tum-l*^’s larval-to-adult viability from 35% to up to 90% (SI Fig. S5E) while 5 mM myristic acid addition had almost no effect (SI Fig. S5E; unlike LA, MA is a fully saturated C18 FA).

Unlike 1 μM and 1 mM aspirin, 5 mM LA treatment did not yield a significant increase in 13-HODE (Fig. 3B), but 13-EFOX-L_2_ was clearly detected in LA-treated *y w* animals and it was 7-fold higher in LA-treated *hop*^*Tum-l*^ mutants than their asprin-treated counterparts (Fig. 3C). These correlations suggest that (a) the double unsaturations in LA are essential for its biochemical transformation to 13-EFOX-L_2_, (b) 13-HODE and 13-EFOX-L_2_ levels can be modulated by dietary modification in the absence of aspirin, and (c) aspirin’s and LA’s effects *in vivo* are realized differently. Overall, however, 13-HODE and 13-EFOX-L_2_ appear to reduce inflammation and improve the survival of *hop*^*Tum-l*^ mutants. We did not detect other members of the EFOX-D or -E families identified in mammalian cells (9), perhaps because they derive from C20 precursors not found in flies (24).

### A blood cell-derived signal affects metainflammation and EFOX levels in drug-free *hop*^*Tum-l*^ animals

STAT92E, the solo STAT family member present in flies, acts downstream of Hopscotch, the only Janus kinase found in flies. Previous reports of *hop*^*Tum-l*^’s cell-autonomous effects (17,33,37) and the tumor-suppressive property of STAT loss-of-function mutations (38,39) suggested that introduction of *msn>STAT*^*RNAi*^ in the *hop*^*Tum-l*^ background should relieve tumor burden. We found that tumor penetrance in *hop*^*Tum-l*^ males is significantly reduced with *msn>STAT*^*RNAi*^ (Fig. 4A), and this reduction may be due, in part, to reduced mitosis in *msn>STAT*^*RNAi*^ macrophages (Fig. 4B). The *msn* promoter is active in many lymph gland progenitors as early as 56 hours (second larval instar (40)) and given that the macrophage/lamellocyte lineages have a shared origin (33,41,42), we hypothesized that mutant blood cells influence systemic health of animals. Targeted knockdown (*msn>STAT*^*RNAi*^ and even *msn>spz*^*RNAi*^) in the overgrowing *hop*^*Tum-l*^ blood cell population rescued metabolic inflammation in larval adipocytes (Fig. 4C, D), while control RNAi did not (SI Fig. S3B, C).

**Fig. 4.**
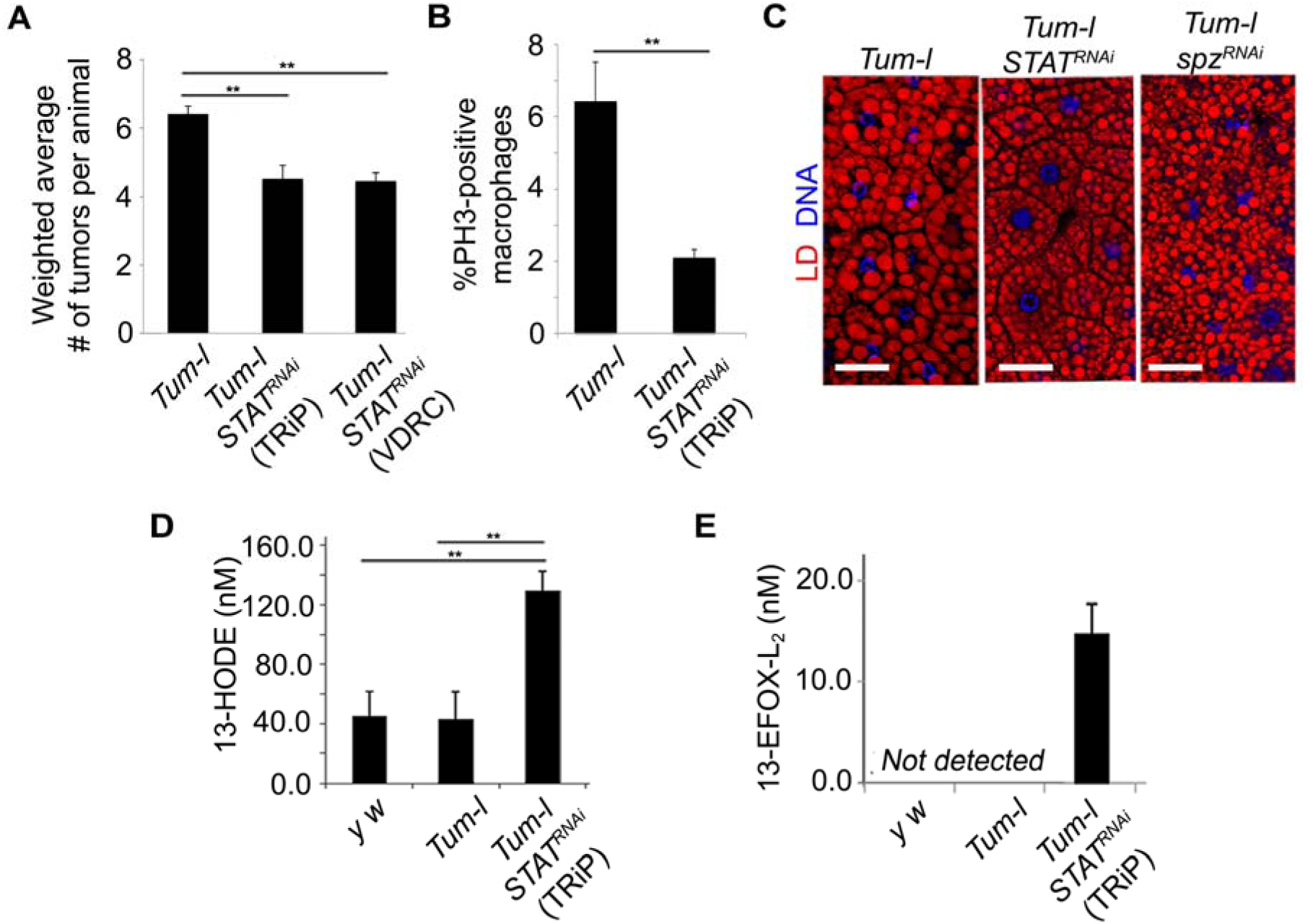
Immuno-metabolic cross-talk: A non-autonomous signal from inflammatory macrophages controls systemic levels of 13-HODE and 13-EFOX-L_2_ in drug-free animals. **(A)** Average number of tumors (weighted for size, see SI) per control (*hop*^*Tum-l*^ *msn>mCD8-GFP*) male or per *hop*^*Tum-l*^ *msn>mCD8-GFP* male expressing *STAT*^*RNAi*^ line (27°C). Pairwise comparisons of each experimental class with control indicate significant differences (*p* < 0.05; student *t*-test). Data from three biological replicates with 28 -52 males in each replicate are represented. The total number of tumors observed ranged from 72 to 189. Error bars are standard errors. The *p* values of unweighted average tumor number comparisons (irrespective of size) were also significant (*p* = 0.02 and 0.03 for the TRiP and VDRC lines, respectively). **(B)** Percentage of PH3-positive macrophages in hemolymph of male larvae is shown. Data were obtained from three biological replicates; hemolymph was examined from 6-10 animals in each replicate. Males were from *hop*^*Tum-l*^ *msn>mCD8-GFP* from *y w* outcross or from cross with TRiP 31318 females. Bars represent averages with standard error; *p*-value is < 0.05; student *t*-test. **(C)** Nile red-stained fat bodies from *hop*^*Tum-l*^ *msn>mCD8-GFP* without or with *STAT*^*RNAi*^ or *spz*^*RNAi*^ expressed in overgrowing blood cells. **(D)** Average number of large, medium, or small LDs per cell from fat bodies of *hop*^*Tum*-l^ mutants as shown. Corresponding images are in Fig. 4C. Bars represent mean ± standard deviation. Three biological replicates were performed and more than 100 cells per replicate were scored. Three animals per replicate were examined; \*\**p* < 0.001, student *t*-test. **(E, F)** 13-HODE and 13-EFOX-L_2_ levels in larval extracts of *hop*^*Tum-l*^ *msn>mCD8-GFP* without or with *STAT*^*RNAi*^ (mean ± standard deviation across four replicates; *n* > 200 larvae per replicate; \*\**p* < 0.05; one-way ANOVA).

This rescue of metabolic inflammation in *hop*^*Tum-l*^ mutants by *STAT* knockdown suggested a correlative rise in anti-inflammatory oxylipids. We observed an increase in 13-HODE (from 43 nM to 130 nM) and 13-EFOX-L_2_ (from undetectable to 14.0 nM) in LC-MS/MS quantification of *hop*^*Tum-l*^ *msn>STAT*^*RNAi*^ animals (Fig. 4E, F). The levels of these oxylipids remained unchanged with control RNAi (SI Fig. S3D, E). These *in vivo* 13-HODE and 13-EFOX-L_2_ levels are consistent with corresponding measurements in mammalian cells (9,36). Together, our observations suggest that a STAT-dependent pro-inflammatory signal from hematopoietic cells affects immune and metabolic health of flies by regulating 13-HODE and 13-EFOX-L_2_ levels.

## Discussion

Basic science and clinical discoveries have revealed the pervasive role of inflammation in human disease. Many of these studies expose the intimate links between diet, drugs, and disease development (2,13-15). Our studies of aspirin’s effects in fruit flies show that aspirin treatment reduces pro-inflammatory cytokine-producing cells and metabolic inflammation. These changes correlate with an increase in anti-inflammatory oxylipids.

Aspirin’s anti-mitotic effects appear to be important to its salutary effects in flies. In both *Ubc9* and *hop*^*Tum-l*^ flies, *spz* expression is high (18,34) and selective knockdown of *spz* in *hop*^*Tum-l*^ blood cells improves organismal viability, much as aspirin treatment does. We envision two possible mechanisms of action: (a) aspirin may block the activities of COX-like enzymes (25,27), which may, in turn, act through the NF-κB/STAT axis in *hop*^*Tum-l*^ blood cells to reduce levels of pro-mitotic/pro-inflammatory Spz protein, or (b) it may block cell cycle progression directly, e.g., by interacting with cell cycle proteins. Aspirin’s anti-mitotic effects and reduction in tumorogenesis in flies are consistent with reported salutary effects in epidemiological studies of cancer patients (43-45). Identifying aspirin’s direct biochemical targets in fly cells (including putative COX enzymes) and understanding its action through these targets will provide new insights into aspirin’s biochemical effects in human cells.

Our biochemical and genetic experiments with the *hop*^*Tum-l*^ larvae revealed the existence of 13-HODE and13-EFOX-L_2_ in flies. Furthermore, these experiments showed that untreated mutants and aspirin-treated mutants differ from the wild type animals in distinct ways: (a) 13-HODE is detected in untreated *y w* controls and *hop*^*Tum-l*^ mutants; its levels are higher in aspirin-treated controls and *hop*^*Tum-l*^ mutants, as well as in *msn>STAT92E* knockdown mutants. (b) 13-EFOX-L_2_ is not detected in untreated control or mutant animals and is found only in aspirin-treated animals or in *msn>STAT*^*RNAi*^ animals. Because lipid droplet morphologies in these mutants with high 13-HODE and13-EFOX-L_2_ are restored, we propose that pro-inflammatory signals arising from blood cells adversely affect the metainflammatory state of the animal. Reducing such signals results in high 13-HODE/13-EFOX-L_2_ levels, reduces metainflammation, and improves viability. This interpretation suggests that aspirin treatment of mutants reverses inflammatory phenotypes by tapping into molecular-genetic pathways that are otherwise inactive in wild type animals. This idea needs to be substantiated further in flies and other organisms, as it potentially provides new avenues for treating inflammatory diseases.

How might 13-HODE and 13-EFOX-L_2_ work in flies? The chemistry of 13-HODE in mammals may suggest some mechanisms to be explored. 13-HODE’s *in vivo* effects in mammals range from modulating pain sensitivity to affecting metabolic syndrome (46). Mammalian 13-HODE serves as a ligand for PPARγ (peroxisome proliferator-activated receptor gamma) (47), whose activation alters gene expression and the distribution of surface cell adhesion molecules (46). For example, in unchallenged human endothelial cells, 13-HODE and the vitronectin receptor are sequestered in the cytoplasm, but upon interleukin-1 stimulation, the receptor is found on the cell surface, where it mediates endothelial/platelet interaction (48). We speculate that differences between adipocyte-macrophage interactions in inflamed and aspirin-treated flies may be mediated by similar changes in cell surface protein composition.

EFOX’s mode of action in mammalian cells appears to be quite different from that of 13-HODE. Half of the cellular EFOX is covalently adducted to proteins, which affects their function. For example, EFOX inhibits the ability of p65, a mammalian NF-κB protein that heterodimerizes with p50 (49), to bind to DNA (9,16). Ectopic activation or ectopic expression of Dorsal and mouse p50 in flies causes hematopoietic tumors of the kind found in *Ubc9* and *hop*^*Tum-l*^ mutants (50,51). These parallel effects of fly and mouse NF-κB proteins suggest that in flies, as in mammals, 13-EFOX-L_2_ can modify protein functions to inactivate immune signaling.

Chronic metabolic and inflammatory disorders together constitute a major health concern worldwide (2, 7-9). The anti-cancer effects of NSAIDs in humans are not yet well-understood (52-54). Aspirin’s anti-mitotic and anti-inflammatory roles parallel its effects in mammals and suggest the existence of molecular similarities underlying recovery from overproliferation, chronic inflammation, and tissue damage in flies and humans. Basic research in *Drosophila* has provided valuable insights in understanding disease development in humans. Our work shows that biochemical and genetic studies in *Drosophila* also have the potential to rapidly clarify genetic mechanisms underlying recovery from inflammatory disease. Such studies can help identify and evaluate therapeutic targets for tumor development, dietary stress, and inflammation at the earliest stages of disease development.

## Materials and Methods

### Fly lines

Stocks with recessive (*y w; Ubc9*^*4-3*^*FRT40A/CyO y*^*+*^, *y w; Ubc9*^*5*^ *FRT40A/CyO y*^*+*^ Ref: (20)) or dominant *y w hop*^*Tum-l*^ (19) mutations were modified by simple crosses to examine cell types or manipulate gene expression by RNAi lines. *Canton S* or the *y w* strain was used as a control. Details of fly strains and assays for examining aspirin’s effects are in SI Materials and Methods.

### Chemicals used and analysis of animals

Aspirin (> 99%), salicylic acid (≥ 99%), linoleic acid (≥ 99%) and myristic acid (≥ 99%) were purchased from Sigma Aldrich (St Louis, MO). Aspirin tablets were from Bayer (325 mg of active principle). Age-matched third instar heterozygous or mutant larvae were chosen from 6-24-hour egg lay without gender bias. Developmentally delayed animals were excluded. Detailed protocols for chemical administration and parameters for (a) tumor penetrance and expressivity, and lymph gland lobe size; (b) mitosis; (c) gene expression; (d) fat body integrity; and (e) Nile red-stained LD measurements in adipocytes are described in the SI Materials and Methods.

### Tumor penetrance and expressivity

Aspirin’s effects on the presence and abundance of tumors (penetrance) and tumor sizes (expressivity) were scored in 8 day-old *Ubc9* animals. In three independent experiments, all available mutant untreated or aspirin-treated animals were first scored for absence/presence of visible tumors using a stereomicroscope. All animals were then dissected and all tumors were mounted on slides and imaged with a Zeiss Axioscope 2 Plus microscope. Surface area measurements (a measure of tumor size) were made using the AxioVision LE 4.5 software. Because of recessive lethality of the mutation, few mutants survive at 8 days.

### Biochemistry

For oxidized lipids in *Drosophila* larvae, we used solid phase extraction methods (55). At least 200 age-matched animals were transferred from fly food onto a fine-sieve mesh, washed with water, 70% ethanol, and then again with water, and gently dried with a Kimwipe. Samples were weighed and stored at -80°C until further use. The details of sample preparation, analysis of lipid derivatives by reverse phase HPLC coupled to a mass spectrometer and parameter for Multiple Reaction Monitoring analyses are explained in SI Materials and Methods.

### Statistical analysis

Experiments were replicated three or more times. Where results were quantified, at least three biological repeats were performed. The numbers of animals tested are indicated in the text, Figure legend, or SI Methods. For pairwise comparison, either one-way ANOVA or the *t*-test was applied. *p* < 0.05 was considered to be significant.

## Acknowledgements

The Mott Hall Middle School students Farrah Lopez and Diego Mendia assisted with initiating this project. Mark Lee, Lawrence Huang, Zoya Huda, and Zoe Papadopol assisted with experiments or data analysis. We thank members of the Govind lab for helpful discussions. We are grateful to R.A. Schulz, Bloomington *Drosophila* Stock Center, and VDRC for fly strains. K. Dastmalchi provided suggestions on lipidomics experiments and L. Yang provided technical support for mass spectrometry. This work was supported by funds from the National Science Foundation (1121817-SG & 1512458-GJ), NASA (NNX15AB42G-SG), and the National Institutes of Health (S06 GM08168, 8G12MD007603-30-RCMI).

## Author contributions

IP, SP, GJ & SG conceived experiments. IP discovered the anti-inflammatory effects of aspirin; SP confirmed aspirin’s effects and discovered the oxidized lipids. Fig. 1: IP, RR, & TG. Fig. 2: RSM, RR, SG & SP. Fig. 3: SP. Fig. 4: JC, SG & SP. All authors contributed to manuscript preparation.

