## Supplementary Information for "A *Drosophila* metainflammation-blood tumor model links aspirin-triggered eicosanoid-like mediators to immune signaling"

### Supplementary Information Text

#### Effects of salicylic acid on *Ubc9* mutants

Salicylic acid (SA) is the principal metabolite of aspirin and lacks the functional acetyl group characteristic of ASA. SA was dissolved in dimethyl sulfoxide (DMSO) to make a 10 M solution. The final DMSO content in the fly food was negligible being four orders of magnitude lower than the reported cytotoxic DMSO concentration in *Drosophila* larvae (~ 0.3% w/w (1)).

SA treatment of *Ubc9* mutants did not ameliorate tumor penetrance of *Ubc9* mutants. At 1 nM and 50 nM concentrations, all animals ($n$ = 10 and $n$ = 13, respectively) developed tumors and produced aggregates, much like the untreated controls. None of the mutants was tumor-free. This result was also reflected in infiltration indices (number of macrophages adhering to adipose tissue) where no significant difference between SA-treated (1.58 ± 0.10 for 1 nM and 2.46 ± 0.27 for 50 nM SA) and control treatment (2.64 ± 0.34, no DMSO, or 2.18 ± 0.06, with DMSO) was observed ($p > 0.05$). In each case, 100 fat body cells were examined for infiltration by macrophages.

No significant difference was observed in tumor penetrance between “no treatment” or “DMSO-only” treatment classes of *Ubc9* animals. Under both conditions, all animals developed tumors and/or aggregates. Infiltration index was also statistically indistinguishable (2.64 ± 0.34 without DMSO and 2.18 ± 0.06 with DMSO; $n = 100$ cells in each case; $p > 0.05$).

**Effects of aspirin on tumor sizes and viability**

Average projection area values for 8-day *Ubc9* tumors from the 0.5 nM aspirin dataset were 181,024 ± 8,571 $\mu m^2$ and 48,144 ± 31,459 $\mu m^2$ for untreated and treated samples, respectively. The corresponding values from the 5.5 µM aspirin dataset for 8-day *Ubc9* animals were 206,308 ± 37,011 $\mu m^2$ and 75,251 ± 11,943 $\mu m^2$ for untreated and treated samples, respectively (± refers to standard deviation; $p < 0.05$).

*Ubc9* tumors are derived from expanded lymph gland lobes (2). A significant reduction in the average projection area between untreated (9 glands, 18 lobes) and 0.5 nM aspirin-treated (6 glands, 12 lobes) anterior or posterior lobes (pairwise comparison, $p < 0.05$, two-tail) was observed. Average size for anterior lobes: untreated 30,462 ± 4,380 $\mu m^2$ versus aspirin-treated 18,164 ± 3,158 $\mu m^2$. Average size for posterior lobes: untreated 20,907 ± 2,467 $\mu m^2$ versus aspirin-treated 9,221 ± 1,662 $\mu m^2$. ± denotes standard error.

Viability of 0.5 nM and 5.5 µM aspirin-treated *y w* animals remained as high as untreated *y w* animals (98-100%) and no toxic effects were observed. Addition of 1 mM aspirin in fly food resulted in partial lethality at pupal stages.

### Materials and Methods

#### Fly lines and crosses

*Ubc9* strains (a) *y w; Ubc9*$^{4-3}$*FRT40A/CyO y*$^{+}$, (b) *y w; Ubc9*$^{5}$ *FRT40A/CyO y*$^{+}$, (c) *y w; Drs-GFP, lwr*$^{4-3}$*/CyO y*$^{+}$, and (d) *y w; Drs-GFP lwr*$^{5}$*/CyO y*$^{+}$ are described in Chiu et al. (2005) (3). The *76B>GFP* inserts in the *Ubc9* background are described in Kalamarz et al. (2012) (2). The X-linked temperature-sensitive *hop*$^{Tum-l}$ mutation (4) was recombined with X-linked *msnf9-GAL4* insert (5). The *mCD8-GFP* reporter transgene (BDSC:5137) was introduced in this background to distinguish macrophages (GFP-negative) from GFP-positive lamellocytes. The final genotype of this stock is: *y w hop*$^{Tum-l}$ *msn-GAL4/FM7; UAS-mCD8-GFP* and is referred to as *hop*$^{Tum-l}$ *msn>mCD8-GFP* for short.

UAS "knockdown" lines are as follows: Control *UAS-RNAi* lines used were: *white* (TRiP lines 25785 or 28980), *ebony* (TRiP line 28612), and *GFP* (TRiP line 41554)). *white* knockdown was validated with the *eyeless-GAL4* driver and *ebony* knockdown was confirmed with universal *daughterless-GAL4 (da-GAL4)* driver. Loss of the red pigmentation in eyes of *ey>w*$^{RNAi}$ flies and darker adult body color in *da>e*$^{RNAi}$ flies was observed. The EGFP.shRNA construct was validated in Neumüller et al. (6).

*UAS-STAT*$^{RNAi}$ lines used for Signal transducer and activator of transcription (STAT92E) were 31318 (TRiP1), 31317 (TRiP2), and VDRC GD 43867. The TRiP 31318 strain was used for biochemistry. The line for *Spätzle processing enzyme* (*SPE*) knockdown was *w; UAS-SPE*$^{RNAi}$ */CyO* (VDRC v30972). For *spätzle* (*spz*) knockdown *y w; UAS-spz*$^{RNAi}$ VDRC v105017 and TRiP 28538 lines were used and for *Toll* (*Tl*) knockdown *UAS-Tl*$^{RNAi}$*/MKRS* TRiP 31044 (TRiP1) and 31477 (TRiP2) were used. *UAS-dl*$^{RNAi}$ TRiP 27650) was used for *dorsal* (*dl*) knockdown,

**Gene silencing in *hop*$^{Tum\text{-}l}$ flies**

Because of the semidominant X-linked mutation, fewer *hop*$^{Tum\text{-}l}$ */Y* males survive relative to their *FM7/Y* siblings. This genetic background is therefore suitable for biochemical and genetic interaction experiments. *y w hop*$^{Tum\text{-}l}$ *msn-GAL4/FM7; UAS-mCD8-GFP* females were crossed either with *UAS-RNAi* males or with *y w/Y* control males. Effects of RNA interference (*STAT*, *SPE*, *spz*, *Toll* or *dl*) on viability in the *y w hop*$^{Tum\text{-}l}$ *msn-GAL4/FM7; UAS-mCD8-GFP* animals were normalized relative to the *y w* outcross. The proportion of *hop*$^{Tum\text{-}l}$*/Y* (mutant) sons relative to *FM7/Y* (control) sons is defined as relative rescue. In additional crosses with control *UAS-RNAi* for *white* (two lines), *ebony*, or *GFP*, we found that in all four pairwise comparisons, the ratio of surviving *hop*$^{Tum\text{-}l}$*/Y* RNAi sons relative to *FM7/Y* balancer-carrying sons was statistically indistinguishable from this ratio derived from the *y w/Y* outcross (Fig. S3A).

**Aspirin purity, administration, and effects on viability**

The sources of aspirin for *Ubc9* and *hop*$^{Tum\text{-}l}$ treatments were different. For *Ubc9* larvae, One Bayer aspirin tablet (325 mg) was ground finely and 1 mg/ml of this powder was dissolved in an appropriate amount of water and the suspension was stirred vigorously for 15 min and filtered. An appropriate volume of aspirin solution was added to fly food to yield aspirin concentrations of 0.5 nM or 5.5 µM. Purity of the drug in these tablets was determined as follows: a 325 mg tablet was ground to a fine powder and suspended in ice-cold methanol with vigorous stirring for 10 min. It was vacuum-filtered with a Buchner funnel. The solvent was dried under a gentle stream of nitrogen; the white solid was further dried under high vacuum, overnight. High purity of the compound (> 99.8%)

was verified via LC-MS and NMR analyses. Results matched data reported in the literature (7).

For *hop*$^{Tum-l}$ larvae, a 10 M aspirin (≥ 99.9% Sigma Aldrich) solution in DMSO was diluted in water to achieve a final aspirin concentration of 1 nM, 1 μM or 1 mM in fly food. The final DMSO content in the fly food was significantly lower than levels known to be toxic in *Drosophila* larvae (~ 0.3% w/w (1)). An appropriate volume of the resulting aqueous solution was added directly to fly food and thoroughly homogenized with an electric mixer for 15 min. Larvae were reared at 27ºC and collected for lipid extraction, viability assays, or dissections, 5 days after egg lay, unless stated otherwise.

Aspiring administration did not rescue recessive lethality of *Ubc9* mutants. To examine aspirin's effects on larval-to-adult viability, the number of *hop*$^{Tum-l}$ *msn>mCD8-GFP* adults emerging from a known number of second/third instar larvae (untreated or treated with 1 nM, 1 μM, or 1 mM of aspirin) was scored.

**Linoleic acid and myristic acid administration**

An appropriate volume of pure LA (≥ 99.9%, Sigma Aldrich, St Louis, MO) was added directly to fly food for a final concentration of 5 mM. The mixture was homogenized into fly food for 15 min. *y w* and *hop*$^{Tum-l}$ *msn>mCD8-GFP* larvae were reared at 27ºC for 5 days and collected for lipid extraction or viability assays. Myristic acid (MA, ≥ 99.9%, Sigma Aldrich) was similarly administered, also at 5 mM final concentration.

**Synthesis of Rhodamine B-conjugated aspirin and visualization**

In a 150 ml two-neck round bottom flask, 2 g (4 mmol) of Rhodamine B was dissolved in 16 ml (10 mmol, 2.5 equivalent) of $POCl_3$ and the mixture was vigorously stirred at 110°C for 2 days. Subsequently, $POCl_3$ was thoroughly distilled off under vacuum after which 60 ml of molecular sieve-dried acetonitrile was added to the flask along with 2.1 g (11.3 mmol, 2.8 equivalent) of mono-Boc-protected piperazine and 0.7 ml (5 mmol, 1.2 equivalent) of triethylamine. The mixture was stirred at room temperature for 24 hours, and then for 12 hours under reflux. Once thin layer chromatography showed complete conversion of the starting material, triethylamine was distilled off and the crude product was dissolved in 20 ml of trifluoroacetic acid and stirred for 2 hours at room temperature. The solvent was removed and the crude product was purified via flash chromatography (initial eluting mixture 5.5 $CHCl_3$: 1 $CH_3OH$ changed gradually to 2:1 ratio). After collecting the correct fractions, the solvent was removed using a rotary evaporator and the final product was precipitated by dissolving it in 2 ml of methanol and then adding it to 50 ml of diethyl ether, drop-wise.

After drying it under vacuum, 150 mg of this intermediate (1 equivalent) was dissolved in 10 ml of DMSO to which 54 mg of aspirin (0.3 mmol, 1 equivalent), 153 μl of triethylamine and 410 mg of HBTU (1 mmol, 3.3 equivalent) were added in this order. The mixture was stirred for 24 hours at room temperature under nitrogen. At the end of this period, 150 ml of 0.5 M $NaHCO_3$ were added followed by 100 ml of ethyl acetate. The biphasic mixture was transferred into a separatory funnel. Upon vigorous stirring the organic phase was recovered, whereas the aqueous phase was extracted 3 times with ethyl acetate. The combined organic fractions were dried over $Na_2SO_4$. After filtration on a cotton plug the solvent was removed by rotary evaporator almost to dryness. The crude product was dissolved in 2 ml of ethyl acetate which were added to 150 ml of diethyl ether drop-wise at room temperature. After keeping the solution at 4°C for 1 hour, the product was recovered via filtration and dried in vacuum to yield 160 mg of Rhodamine B-conjugated aspirin (RhASA, 80% yield over two steps; 1H-NMR: 400 MHz, δ (ppm): 1.13 (12H, t), 2.08 (3H, s), 3.39 (12H, q)), 3.50 (4H, t), 3.60 (4H, t), 5.34 (1H, d), 6.13 (2H, s), 6.19 (2H, d), 6.84 (2H, dd), 7.24 (1H, s), 7.25 (3H), 7.39 (1H), 7.48 (1H, dd), 7.83 (1H, d), 7.92 (1H, d); MS: $[M\text{-}Cl^{-}]^{+}$ = 444.24 m/z). A working solution (800 μM RhASA in phosphate buffer saline (PBS, pH 7.2) was applied to dissected larval tissues for 30 min at room temperature in a humidified chamber. Negative control samples were exposed to a mixture of 800 μM Rhodamine B + 800 μM aspirin solution in PBS.

**Analysis of aspirin-treated animals**

Age-matched third instar heterozygous or mutant larvae were chosen without bias to sex from a timed (6-24 hour) egg lay. Developmentally delayed larvae (small in size) were not selected for analysis.

**(a) Dissection and immunostaining** Fat bodies, lymph glands, blood cell smears, or tumors were dissected from *y w*, heterozygous or mutant larvae (2,8). Air-dried blood cells, aggregates, and tumor samples were stained according to protocols described in (8,9). Samples were mounted in either 70-80% glycerol or in Vectashield (Vector Labs) and were imaged with a Zeiss 510 or 710 laser scanning confocal microscope and formatted in Zeiss LSM5 or Zen 2.3 software.

**(b) Tumor and lymph gland lobe size**

Larval tumors: For frequency and size of blood tumors, age-matched *Ubc9* larvae were collected 8 days after egg lay. Washed animals were bled; hemolymph contents were recovered on glass slides and air-dried, fixed in 4% paraformaldehyde in PBS (pH 7.2), mounted in 80% glycerol and photographed using a CCD camera of a Zeiss Axioscope. AxioVision LE Rel. 4.5 software was used to measure projection areas. This term refers to the area internal to the outline of a fixed, mounted, and imaged structure. Structures smaller than 10,000 $\mu m^2$ in projection area were recorded as aggregates and are not reported, while those larger than 10,000 $\mu m^2$ were defined as microtumors or tumors (2).

Lymph gland lobe size: The projection areas of individual anterior and first set of larva posterior lobes (of the same gland) from age-matched, untreated or 0.5 nM aspirin-treated 6-day old larvae were measured in the same way as tumors and aggregates above.

Tumors in adult flies: All tumors in dorsal and ventral abdominal areas were scored for (i) *hop$^{Tum-l}$ msn>mCD8-GFP* males (from a cross of *hop$^{Tum-l}$ msn-GAL4/FM7; UAS-mCD8-GFP* females with *y w/Y* males) and (ii) *hop$^{Tum-l}$ msn-GAL4/Y; UAS-mCD8-GFP* males, expressing *STAT$^{RNAi}$* (lines VDRC GD 43867 or TRiP line 31318), cultured at 27°C. A tumor was considered small if it was less than half the length of the body segment, medium-sized if it spanned more than half but less than a complete body segment, and large if the tumor's surface area exceeded one entire segment. A medium and large tumor was weighted to equal 2 or 3 small tumors, respectively. The average number of tumors (weighted for size in this way) was computed for each cross and pairwise comparisons of average number of tumors/animal in each experimental class with control class were made to determine differences due to *STAT*$^{RNAi}$ expression. Larval hemolymph was examined for mitotic index (see below).

**(c) Mitosis** For *hop*$^{Tum-l}$ *msn>mCD8-GFP* blood cells (without or with aspirin), the rabbit anti-PH3 (1:200; EMD Millipore) was visualized with anti-rabbit alkaline phosphatase-linked secondary antibody (1:5000; Thermo Scientific). Alkaline phosphatase staining was performed with 125 μg/ml BCIP and 250 μg/ml NBT from Promega. Mitotic indices of control *hop*$^{Tum-l}$ *msn>mCD8-GFP* larvae from *y w* cross and *hop*$^{Tum-l}$ *msn>mCD8-GFP* animals expressing *STAT*$^{RNAi}$ (TRiP 31318) was determined using the same methods, except hemolymph from only male animals was examined in each case.

**(d) Reporter gene expression** Changes in *Drs-GFP* expression in third instar larval fat bodies, 6 days and 8 days after egg lay were assessed. The entire fat body was carefully dissected and mounted on a slide. The status of GFP expression in the organ was scored under a Zeiss Axioplan 2. AxioVision LE 4.5 software was used to process images.

**(e) Infiltration index** Third instar larval fat bodies were dissected from untreated control and aspirin-treated larvae. Dissected tissues were fixed and stained with Hoechst 33258 and TRITC-labelled Phalloidin. Samples were mounted in 50% glycerol in PBS, 7.4. Using Zeiss Axioscope fluorescence microscope, the number of single blood cells adhering above or below the fat body were scored as described in (9).

**(f) Slide preparation for fat body lipid droplet analysis** Fat body LDs were detected after fixation and staining with Nile red (N3013, Sigma Aldrich) as described by Greenspan et al. (10). Samples were kept in PBS (pH 7.2) throughout the procedure. Samples were counterstained with nuclear dye Hoechst 33258 (both from Invitrogen Molecular Probes) and were mounted in Vectashield. To preserve the endogenous LD sizes, slides carrying fat body samples were prepared as follows: Fixed, Nile red- and Hoechst 33258-stained fat body pieces were mounted on slides with a spacer to prevent distortion of LDs and maintain normal tissue morphology. (i) A 4-5-cm strip of single-sided Scotch tape was placed in the center of the slide. (ii) A thin, even layer of Vaseline was applied onto the tape with a paintbrush. (iii) Using the corner of a razor blade, a 1 cm x 3 cm rectangle was excised from the tape and peeled off using tweezers. (The exposed rectangular area becomes the well for the sample.) (iv) A drop of Vectashield was placed in the well and evenly spread on the glass surface. (v) Using clean tweezers, 3-4 pieces of the stained fat bodies (in PBS, pH 7.4) were transferred into the well avoiding overlap. (vi) A glass cover slip was then gently placed over the samples and uniformly pressed; the Vaseline adheres the tape to the coverslip. The sample in the Vectashield should now occupy the well. (vii) Clear nail polish was gently applied to seal the coverslip. While the prepared slides can be stored at 4°C, optimal results were obtained when samples were imaged immediately. Samples were imaged on a Zeiss LSM 510 or 710 confocal microscope. Lipid droplet sizes were classified as follows: small (radius, $r < 3.1$ μm); medium ($3.1 < r < 5$ μm); large ($r > 5$ μm).

**Biochemistry experiments**

**(a) Materials for lipidomics** Solid-phase extraction STRATA SPE cartridges: C18-E (500 mg, 6 ml; Cat. No. 8B-S001-HCH); non-chiral RP-HPLC column: Phenomenex Luna 3 μm C18(2) 100A 150x2.0 mm; Chiral HPLC column: Chiracel OD-RH 5 μm 150x2.1 mm; MS system: 4000 QTRAP (Applied Biosystems), HPLC system: Shimadzu Prominence HPLC (Shimadzu USA); NMR instrument: Varian Mercury-300. MS-grade water, acetonitrile and formic acid were purchased from WorldWide Life Sciences (Bristol, PA). The following analytical-grade standards were purchased from Cayman Chemical Co. (Ann Arbor, MI): 13(S)-HODE (>98%), 13-EFOX-$L_2$ (>98%), 13(S)-HOTrE (>98%), 13(S)-HODE-$d_4$ (>99% deuterated), 13-EFOX-$L_2$-$d_3$ (>99% deuterated).

**(b) Sample preparation** The amounts of solvents and internal standards before solid phase extraction used for 200 mg of larvae are presented below; these amounts were adjusted where larval weight exceeded 200 mg. For lipid extraction, larvae were transferred into a 1 ml pre-chilled glass Dounce grinder and homogenized thoroughly at 0ºC with a glass pestle along with 500 µl of cold methanol. The pestle was rinsed twice with 100 µl of cold methanol. 4 ml of ice-cold water were added to obtain a 15% methanol solution. Subsequently, 50 ng of deuterated internal standards were added to the homogenate which was left on ice in the dark. After 15 min the homogenate was centrifuged at 4ºC and 4,000 rpm for 10 min. The supernatant was recovered and kept on ice. The pH was then adjusted to 3 by adding 10-20 drops of 0.1 M HCl and 2.00 ml of the acidic extract were immediately loaded onto the SPE column which was previously conditioned with 20 ml of methanol and 20 ml of water. The cartridge was washed with 20 ml of ice-cold 15% methanol in water, 20 ml of water and 10 ml of hexane. The lipids were recovered with 12 ml of cold methyl formate which was evaporated on ice and in the dark under a gentle stream of nitrogen. The purified lipids were dissolved in 1 ml of cold, degassed ethanol, and stored at -80ºC until the LC-MS analysis.

Lipid derivatives were analyzed on a reverse phase-high pressure liquid chromatography coupled to a mass spectrometer (RP-HPLC MS/MS). The non-chiral LC-MS runs were performed after diluting the extract ten-fold in ethanol. All samples were analyzed with a gradient solvent system consisting of A (water with 0.1% formic acid) and B (acetonitrile with 0.1% formic acid). The flow rate was 250 µl/min and the gradient used was the following: hold at 35% B for 3 min, then 35-90% B in 23 min, then 90-100% B in 0.1 min, hold for 5.9 min and 100-35% B in 0.1 min, and finally hold for 7.9 min. The oven temperature was 40ºC. Chiral analyses were performed in isocratic elution with 35% A and 65% B for 25 min. The flow rate was 250 µl/min and the oven temperature was 40ºC.

LC-MS analyses were performed in Multiple Reaction Monitoring (MRM) mode whose optimal parameters and most abundant fragments were obtained experimentally for 13-HOTrE, 13-HODE, and 13-EFOX-$L_2$ by utilizing commercially available synthetic standards. The settings used are shown in Table 1. Standard curves were prepared using 13-HODE, and 13-EFOX-$L_2$ as analytes and 13-HODE-$d_4$ and 13-EFOX-$L_2$-$d_3$ as internal standards. Analyte/internal standard peak area ratios were used for quantification. Comparison of retention times and the use of at least two MS transitions per compound rendered the detection reliable and sensitive (Fig. S5B, C and Fig. S6). Data were analyzed by one-way ANOVA.

**Figure S1**

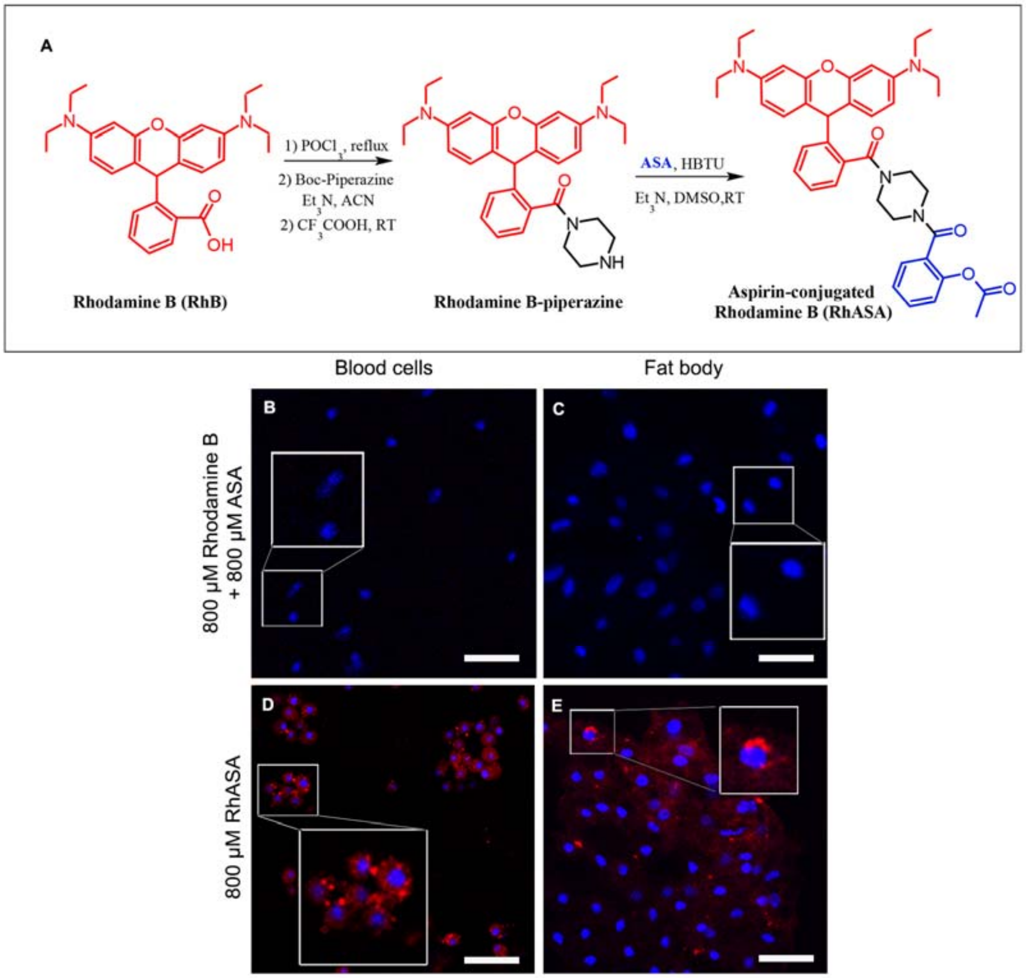


**Fig. S1. Aspirin uptake by fly cells. (A)** Synthesis of Rhodamine B-labeled aspirin. Rhodamine B was first mono-amidated with piperazine. Then, the carboxylic acid moiety of aspirin was conjugated to the secondary amine in the piperazine moiety. RT refers to room temperature.

**(B-E)** Incubation of live *Ubc9*$^{-/-}$ larval tissues with unconjugated Rh and free aspirin (B, C), or 800 µM aspirin-conjugated Rhodamine B (RhASA; D, E). Uptake in circulating blood cells (B, D) and fat body adipocytes (C, E) is shown. Unconjugated Rh is not retained but subcellular localization of RhASA is clearly observed (C and E). Cells were counterstained with Hoechst 33258 after fixation. Scale bars show 15 µm (B & D), and 50 µm (C & E). Cells in the larger box (panels D and E) were zoomed in for clarity.

**Figure S2**

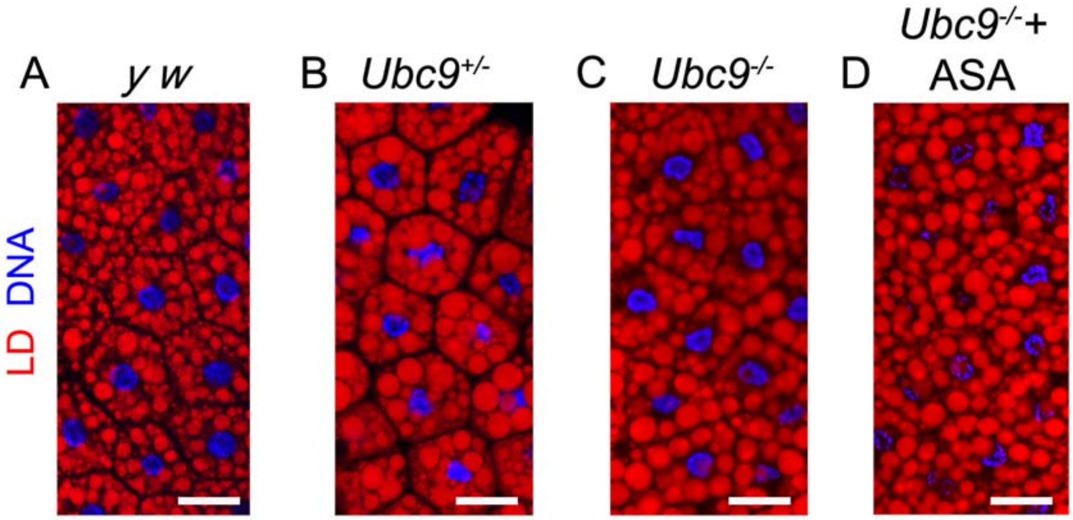


**Fig. S2. Aspirin administration attenuates aberrant lipid droplet morphology.** Metainflammation in *Ubc9* fat body. Nile red-stained LDs in the control (*y w*) background are small and regularly spaced around the nucleus. LDs in *Ubc9* heterozygous or mutant *(*$Ubc9^{4\text{-}3/5}$) adipocytes are larger relative to LDs from *y w* adipocytes. 1 µM aspirin treatment partially rescues metainflammation. Scale bar = 30 µm.

**Figure S3**

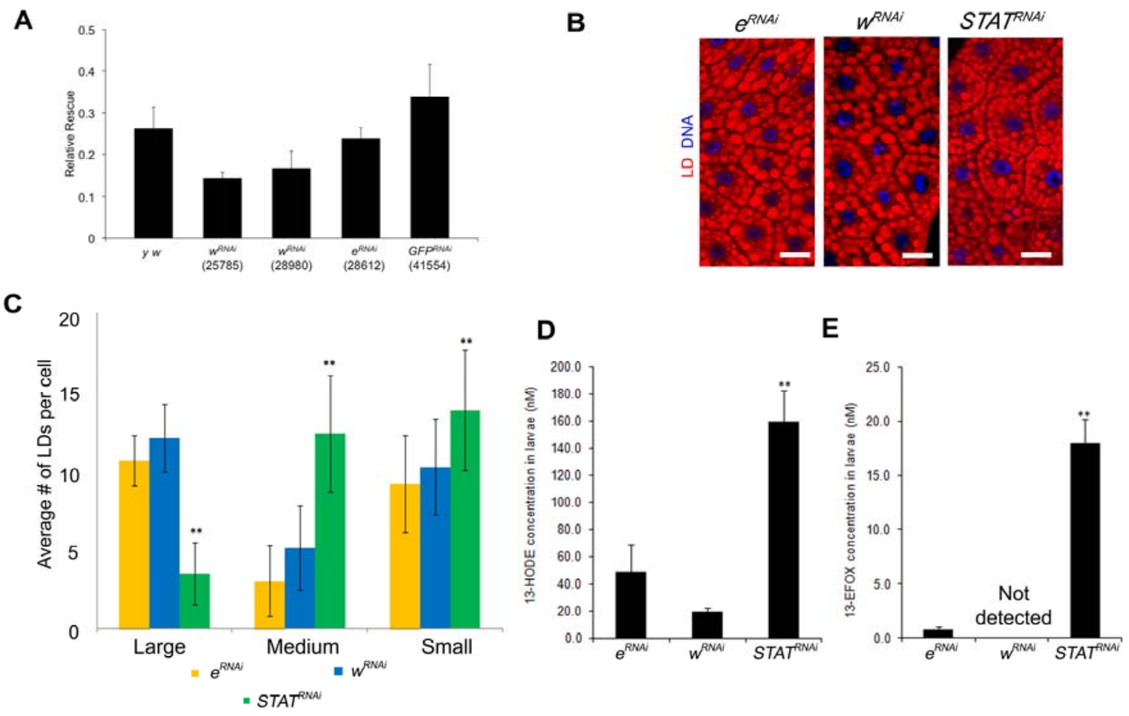


**Fig. S3. Control experiments: Gene silencing.**

**(A)** Viability of control *hop*$^{Tum\text{-}l}$ *msn/Y; UAS-mCD8-GFP/+* males (born from a cross of *hop*$^{Tum\text{-}l}$ *msn-GAL4/FM7; UAS-mCD8-GFP* females with *y w/Y* males) compared to *hop*$^{Tum\text{-}l}$ *msn-GAL4/Y; UAS-mCD8-GFP/+* males expressing one copy of RNAi of *white*, *ebony*, or *GFP* raised at 27°C. The proportion of observed mutant males relative to *FM7/Y* balancer class is shown. Pairwise comparison of the proportion of surviving *hop*$^{Tum\text{-}l}$ males expressing RNAi with the *y w* outcross shows no significant difference ($p > 0.05$; *p*-values ranged from 0.10 to 0.71, student *t*-test; mean ± standard error). Three biological replicates with at least 100 males in each replicate are represented. Error bars are standard errors. See SI Methods for details.

**(B, C)** Nile red-stained fat bodies (B) and average number of LD size categories per fat body cell (C) from progeny of homozygous *hop*$^{Tum\text{-}l}$ females crossed with *y w* males or males homozygous for UAS-*white*$^{RNAi}$, *ebony*$^{RNAi}$, or *STAT*$^{RNAi}$ transgenes. Bars represent mean ± standard deviation computed across three biological replicates; $n > 100$ cells, 6 animals each; $^{**}p < 0.001$, student *t*-test.

**(D, E)** 13-HODE and 13-EFOX-$L_2$ levels in larval extracts of animals with genotypes in panel B and C (mean ± standard deviation; $n > 200$ larvae per replicate; $^{**}p < 0.05$ across three biological replicates, one-way ANOVA).

**Figure S4**

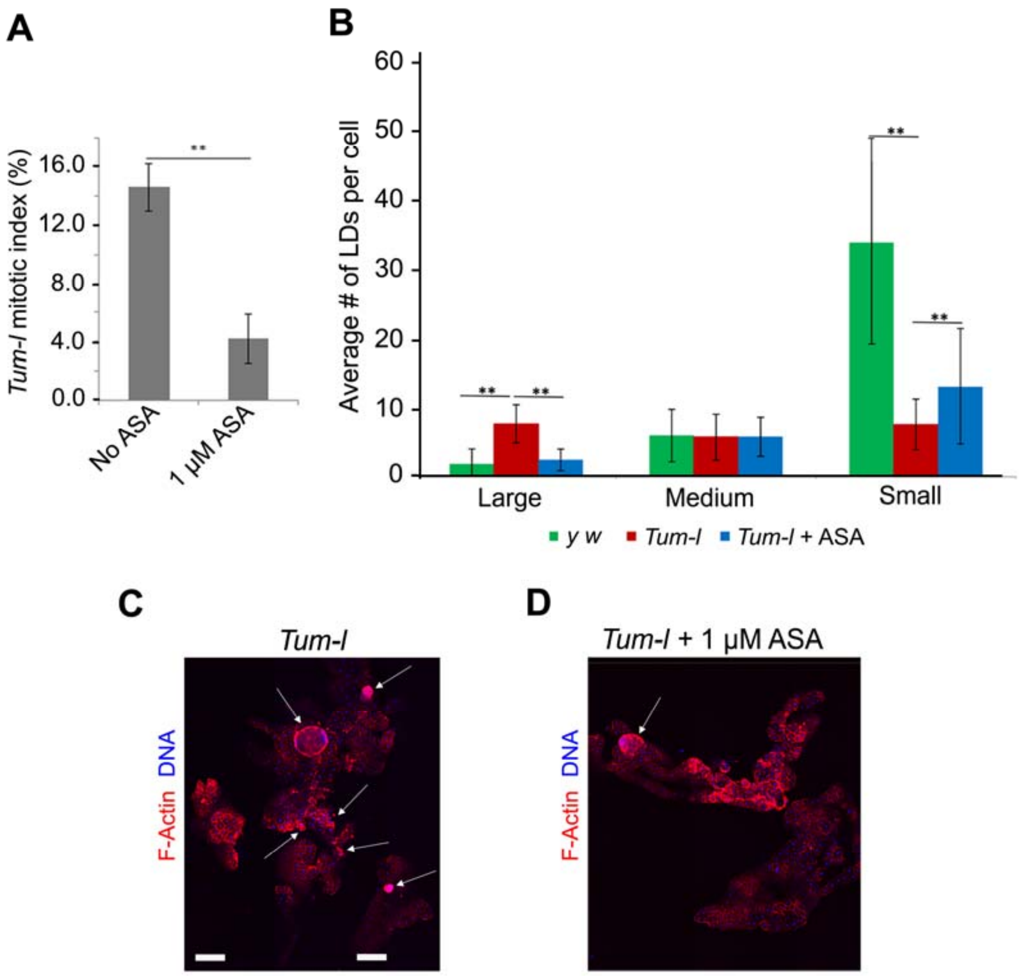


**Fig. S4. Effects of aspirin treatment on *hop$^{Tum-l}$* animals.**

**(A)** Mitotic index in blood cell smears of *hop$^{Tum-l}$ msn>mCD8-GFP* animals upon systemic administration of 1 μM aspirin (mean ± standard deviation; three biological replicates, 20 larvae per replicate, and more than 800 macrophages per treatment; ** $p < 0.01$, student *t*-test).

**(B)** Average number of large, medium, or small LDs per cell from fat bodies of untreated *y w* animals, and untreated and 1 μM aspirin-treated *hop$^{Tum-l}$ msn>mCD8-GFP* animals. Corresponding images are in Fig. 2F. At least 30 fat body cells per animal were scored and mean ± standard deviation was computed using 6 animals. Three biological replicates were performed. (**$p < 0.001$, student *t*-test.)

**(C, D)** Confocal images of Rhodamine-Phalloidin and Hoechst 33258-stained fat body from untreated (C) or 1 μM aspirin treated (D) *hop$^{Tum-l}$ msn>mCD8-GFP* animals. Fewer aggregates and tumors were present on the organ from treated animals and the organ morphology is closer to normal.

**Figure S5**

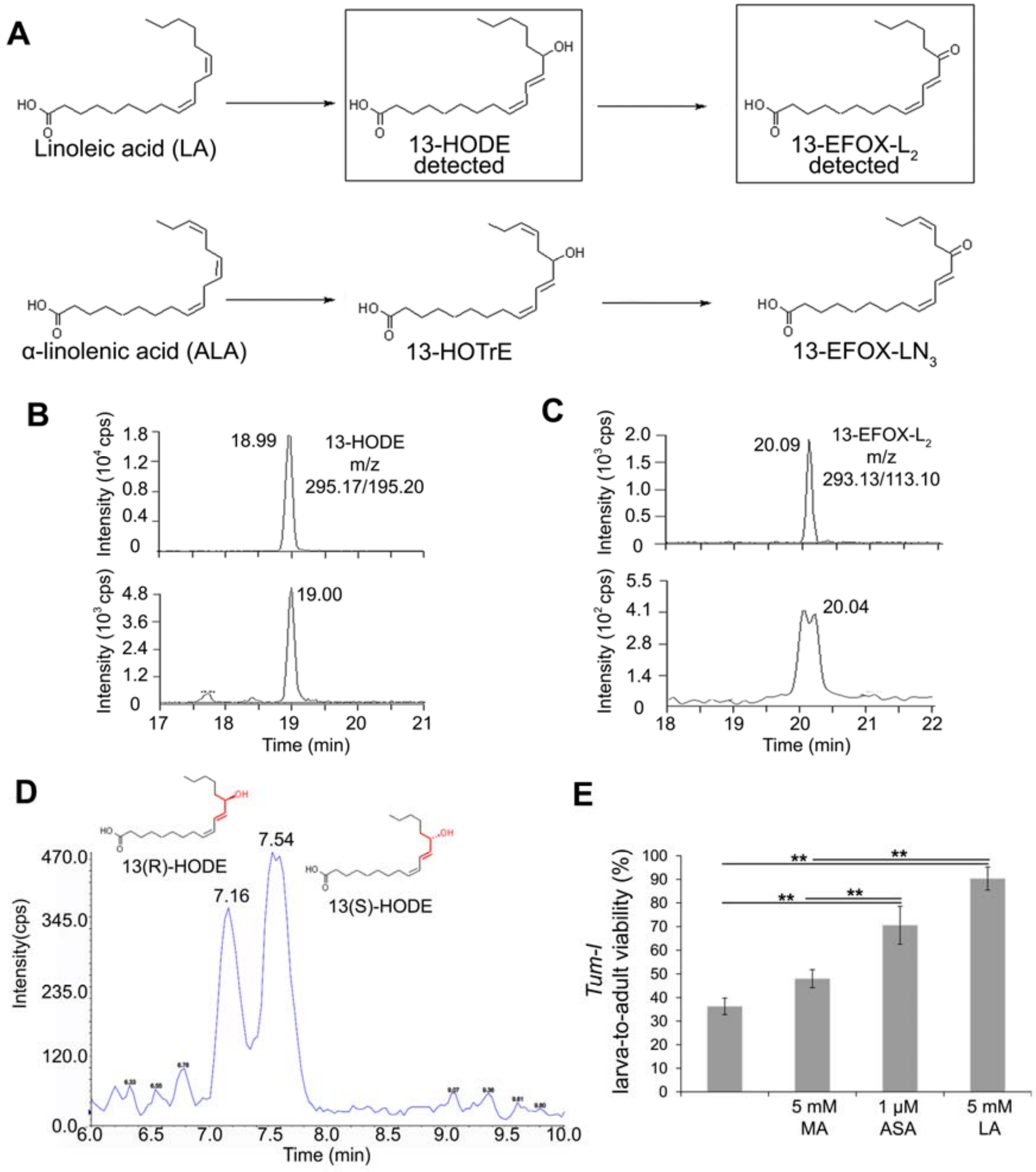


**Fig. S5. Lipidomics analysis and larva-to-adult viability.**

**(A)** Proposed biochemical mechanism based on the scheme by Groeger et al. (11) LA and ALA conversion to 13-HODE and 13-HOTrE, respectively by COX-2 action. These are then further oxidized to 13-EFOX-L2 and 13-EFOX-LN3, respectively, via dehydrogenase enzymes.

**(B)** MRM scans monitoring for the m/z transitions 295.17/195.20 (MW of 13-HODE/loss of 99.97 m/z, see Fig. S5A) for standard 13-HODE (top panel) and a representative lipid sample (bottom panel).

**(C)** MRM scans monitoring for the m/z transitions 293.13/113.10 (MW of 13-EFOX-$L_2$ /loss of 180.03 m/z, see Fig. S5B) for standard 13-EFOX-$L_2$ (top panel), and a representative lipid sample (bottom panel).
**(D)** Chiral LC-MS analysis of larval extracts from 1 μM aspirin-treated *hop*$^{Tum\text{-}l}$ animals. Representative chiral chromatogram showing two-fold enantiomeric excess of 13(S)-HODE observed in larval extracts of wild type and mutant animals. This enantiomeric excess of 13(S)-HODE was also observed in aspirin-treated animals.
**(E)** Effect of myristic acid (MA), aspirin or linoleic acid (LA) treatment on viability of *hop*$^{Tum\text{-}l}$ *msn>mCD8-GFP* larvae. Effects of LA are slightly stronger than those of 1 μM aspirin, while MA's effects are not significant. The proportion of larvae that survived to adulthood was determined in three biological replicates; the number of larvae examined in each replicate ranged from 38 to 68 (mean ± standard deviation; student *t*-test). Statistically significant differences ($p < 0.05$; student *t*-test) are shown across three or four biological replicates.

**Figure S6**

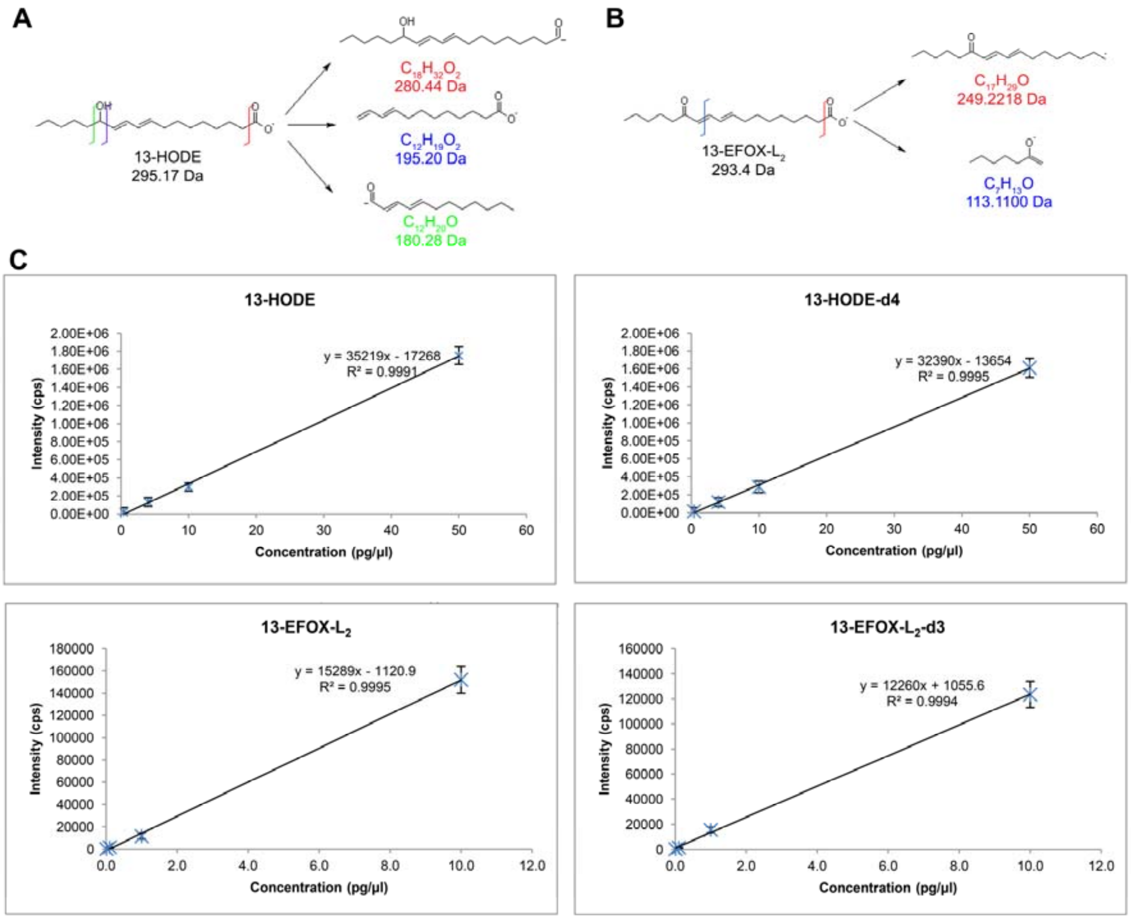


**Fig. S6. Fragments selected for MS/MS analysis and calibration curves.**
**(A, B)** Detection of 13-HODE (A) and 13-EFOX-$L_2$ (B) in MRM mode was based on the structures of the fragments shown. Such fragments were chosen experimentally after ascertaining that they represented the ones forming with the highest yield during the MS/MS analysis.
**(C)** Calibration curves for quantification of the bioactive lipids. The concentrations used to make the calibration curves for 13-HODE-$d_4$ and 13-HODE are 0.4, 4, 10, and 50 pg/µl whereas 0.01, 0.1, 1, and 10 pg/µl concentrations were used to prepare calibration curves for 13-EFOX-$L_2$-$d_3$ and 13-EFOX-$L_2$. The experimental samples were diluted so that analyte concentration was within the linear range (mean ± standard deviation, $n = 6$ samples per analyte per concentration).

**Table S1. Mass spectrometry parameters for Multiple Reaction Monitoring.**

| Compound | Q1 Mass (Da) | Q2 Mass (Da) | DP (Volts) | CE (Volts) |
|---|---|---|---|---|
| 13-HODE | 295.170 | 280.400 | -90 | -26 |
| | 295.170 | 195.200 | -90 | -26 |
| | 295.170 | 180.300 | -90 | -28 |
| 13-EFOX-$L_2$ | 293.130 | 249.200 | -90 | -26 |
| | 293.130 | 113.100 | -100 | -28 |
| 13-HODE-d4 | 299.170 | 281.200 | -95 | -26 |
| | 299.170 | 230.800 | -95 | -16 |
| | 299.170 | 198.100 | -95 | -26 |
| 13-EFOX-$L_2$-d3 | 296.230 | 252.200 | -90 | -26 |
| | 296.230 | 114.100 | -95 | -29 |
